# Basal forebrain GABAergic innervation of olfactory bulb periglomerular interneurons

**DOI:** 10.1101/216259

**Authors:** Alvaro Sanz Diez, Marion Najac, Didier De Saint Jan

## Abstract

Olfactory bulb circuits are dominated by multiple inhibitory pathways that finely tune the activity of mitral and tufted cells, the principal neurons, and regulate odor discrimination. Granule cells mediate interglomerular lateral inhibition between mitral and tufted cells lateral dendrites whereas diverse subtypes of periglomerular (PG) cells mediate intraglomerular lateral inhibition between their apical dendrites. Deep short axon cells form broad intrabulbar inhibitory circuits that regulate both populations of interneurons. Little is known about the extrabulbar GABAergic circuits that control the activity of these various interneurons. We examined this question using patch-clamp recordings and optogenetics in olfactory bulb slices from transgenic mice. We show that axonal projections emanating from diverse basal forebrain GABAergic neurons densely project in all layers of the olfactory bulb. These long-range GABAergic projections provide a prominent synaptic input on granule and short axon cells in deep layers as well as on selective subtypes of PG cells. Specifically, three different subclasses of type 2 PG cells receive robust and target-specific basal forebrain inputs but have little local interactions with other PG cells. In contrast, type 1 PG cells are not innervated by basal forebrain fibers but do interact with other PG cells. Thus, attention-regulated basal forebrain inputs regulate inhibition in all layers of the olfactory bulb with a previously overlooked synaptic complexity that further defines interneuron subclasses.

**Key points summary:**

- Basal forebrain long-range projections to the olfactory bulb are important for olfactory sensitivity and odor discrimination.
- Using optogenetics, we confirm that basal forebrain afferents mediate IPSCs on granule and deep short axon cells. We also show that they selectively innervate specific subtypes of periglomerular (PG) cells.
- Three different subtypes of type 2 PG cells receive GABAergic IPSCs from the basal forebrain but not from other PG cells.
- Type 1 PG cells, in contrast, do not receive inputs from the basal forebrain but do receive inhibition from other PG cells.
- These results bring new light on the complexity and specificity of glomerular inhibitory circuits, as well as on their modulation by the basal forebrain.

## Introduction

Olfactory bulb circuits transform the spatially organized sensory input carried by olfactory sensory neurons (OSNs) into a temporal code in mitral and tufted cells, the principal neurons that relay the information to cortical areas. In the olfactory bulb, inhibitory interneurons vastly outnumber mitral and tufted cells and form several inhibitory circuits that shape the output of principal neurons. In addition, thousands of newborn inhibitory neurons are integrated every postnatal day in the pre-existing olfactory bulb network. Inhibition thus plays fundamental roles in early olfactory processing, as demonstrated by multiple evidence *in vivo* (Abraham *et al*., 2010; Lepousez & Lledo, 2013; Fukunaga *et al*., 2014; Gschwend *et al*., 2015; Economo *et al*., 2016).

Olfactory bulb interneurons are diverse but three large classes dominate: granule cells, periglomerular (PG) cells and short-axon (SA) cells. Granule cells, the most abundant type, form reciprocal dendrodendritic synapses with the lateral dendrites of mitral and tufted cells in the external plexiform layer. PG interneurons, in contrast, interact with the apical dendrites of mitral and tufted cells within the glomeruli. They constitute >80% of the juxtaglomerular GABAergic cells, usually project within a single glomerulus, often lack an axon and are functionally and chemically highly diverse (McQuiston & Katz, 2001; Kosaka & Kosaka, 2007; Panzanelli *et al*., 2007; Parrish-Aungst *et al*., 2007; Whitman & Greer, 2007; Kosaka & Kosaka, 2011; Najac *et al*., 2015; Benito *et al*., 2018). They have been classified in two broad functional classes depending on their excitatory inputs. The majority are so-called type 2 PG cells and receive excitatory inputs exclusively from the dendrites of mitral and tufted cells whereas type 1 PG cells receive direct excitatory inputs from OSNs (Kosaka & Kosaka, 2007; Shao *et al*., 2009). This simple synaptic classification, however, does not fully reflect the functional heterogeneity of PG cells (McQuiston & Katz, 2001; Najac *et al*., 2015; Benito *et al*., 2018). Finally, SA cells with soma located in the glomerular layer (superficial SA cells) or in the granule cell layer (deep or dSA cells) form broad intrabulbar connections. Diverse subtypes of deep SA cells mediate a widespread interneuron-specific GABAergic inhibition (Eyre *et al*., 2008; Boyd *et al*., 2012; Burton *et al*., 2017) whereas superficial SA cells are tyrosine hydroxylase (TH)-expressing mixed dopaminergic/GABAergic neurons that regulate glutamate release from OSNs and monosynaptically inhibit external tufted cells (Liu *et al*., 2013; Whitesell *et al*., 2013).

Granule, PG and SA cells receive GABAergic synaptic inputs but only a few studies have investigated their origin. These inputs could arise from local intrabulbar interactions or from extrabulbar afferences. The olfactory bulb receives centrifugal GABAergic projections from the basal forebrain, particularly from the horizontal limb of the Diagonal Band of Broca (HDB) and from the magnocellular preoptic nucleus (MCPO) (Zaborszky *et al*., 1986; Gracia-Llanes *et al*., 2010; Niedworok *et al*., 2012). These long-range projections make functional connections with granule (Nunez-Parra *et al*., 2013) and dSA cells (Case *et al*., 2017) and their pharmacological inactivation impairs olfactory sensitivity (Nunez-Parra *et al*., 2013). Little is known, however, about their connections in the glomerular layer. We here examined the targets of centrifugal basal forebrain axons in olfactory bulb interneurons. We used patch-clamp recording and light activation of channelrhodopsin-2 (ChR2) targeted to basal forebrain GABAergic neurons in acute slices from dlx5/6-Cre transgenic mice. We specifically focused on PG cells and examined whether GABAergic inputs differ in distinct subclasses of PG cells. The results indicate that long-range basal forebrain GABAergic projections broadly regulate olfactory bulb circuits and innervate specific PG cell subtypes with a previously unexpected level of synaptic complexity.

## Materials and Methods

### Animals

All experimental procedures were approved by the French Ministry and the local ethic committee for animal experimentation (C.R.E.M.E.A.S). We used dlx5/6-Cre female mice that express the Cre recombinase under the control of the regulatory sequences of the dlx5 and dlx6 genes (Monory *et al*., 2006), and Kv3.1-EYFP mice of either sex that express the enhanced yellow fluorescent protein (EYFP) under the control of the Kv3.1 K^+^ channel promoter (Metzger *et al*., 2002).

### Slice preparation

3 weeks to 3 month-old mice were killed by decapitation and the olfactory bulbs rapidly removed in ice-cold oxygenated (95% O2-5% CO2) solution containing (in mM): 83 NaCl, 26.2 NaHCO3, 1 NaH2PO4, 2.5 KCl, 3.3 MgSO4, 0.5 CaCl2, 70 sucrose and 22 D-glucose (pH 7.3, osmolarity 300 mOsm/l). Horizontal olfactory bulb slices (300 µm) were cut using a Microm HM 650V vibratome (Microm, Germany) in the same solution; incubated for 30-40 minutes at 34°C; stored at room temperature in a regular artificial cerebrospinal fluid (ACSF) until use. ACSF contained (in mM): 125 NaCl, 25 NaHCO3, 2.5 KCl, 1.25 NaH2PO4, 1 MgCl2, 2 CaCl2 and 25 D- glucose and was continuously bubbled with 95% O2-5% CO2.

### Electrophysiological recordings

Experiments were conducted at 32-34°C under an upright microscope (SliceScope, Scientifica, Uckfield, UK) with differential interference contrast (DIC) optics. Whole-cell recordings were made with glass pipettes (3-6 MΩ) filled with a K-gluconate-based internal solution containing (in mM): 135 K-gluconate, 2 MgCl_2_, 0.025 CaCl_2_, 1 EGTA, 4 Na-ATP, 0.5 Na-GTP, 10 Hepes (pH 7.3, 280 mOsm). Atto 594 (10 µM, Sigma) was added to the internal solution in order to visualize the cell morphology during the recording. Loose cell-attached recordings (15-100 MΩ seal resistance) were made with pipettes filled with ACSF. Olfactory nerves (ON) projecting inside a given glomerulus were electrically stimulated using a theta pipette filled with ACSF as previously described (Najac *et al*., 2015). The electrical stimulus (100 µs) was delivered using a Digitimer DS3 (Digitimer, Welwyn Garden City, UK). Optical stimulation was done through a blue (490 nm) CoolLED pE 100 (CoolLED Ltd., Andover, UK) directed through the 40X objective of the microscope.

### Cell selection

Visual online inspection of dye-filled neurons combined with functional properties were used to select the cells included in our analysis. Morphology was assessed during recording upon visual inspection of dye-filled cells. PG cells were selected based on the small size (5-7 µm) of their cell body, their position close to the edge of the glomerulus and their small dendritic tree that ramified in a single glomerulus. Superficial SA cells, with larger cell bodies and processes that broadly ramified between or within several glomeruli were excluded from the analysis. PG cell morphology was not reconstructed post-hoc because pipette withdraw after recording inevitably damaged these cells. Deep SA cells were most often found within the IPL, sometimes within the granule cell layer and selected based upon their cell body that was larger (>10 µm) than granule cells’ soma. Moreover, many had a spontaneous high-frequency firing in the cell-attached mode (Eyre *et al*., 2008). Cell morphology was reconstructed in a subset of dSA and granule cells. To do so, neurobiotin (Vector Laboratories INC., Burlingame, CA) was added to the intracellular solution (1 mg/ml). The patch pipette was slowly withdrawn after the recording to avoid damaging the cell body. The slice was then fixed in 4% paraformaldehyde overnight, washed 3 times with PBS and incubated in a permeabilizing solution containing Cy-5 conjugated streptavidin (1 µg/ml; Thermo Fischer Scientific, Waltham, MA) overnight. After 3 rinses with PBS, sections were mounted. Labeled cells were imaged with a confocal microscope (Leica TCS SP5, Leica Microsystems, Wetzlar, Germany).

Recordings were acquired with a multiclamp 700B amplifier (Molecular Devices, Sunnyvale, CA), low-passed filtered at 2-4 kHz and digitized at 10 kHz using the AxoGraph X software (Axograph Scientific). In current-clamp recordings, a constant hyperpolarizing current was injected in order to maintain the cell at a potential of −60/-70 mV. In voltage-clamp recordings, access resistance (Ra<30 MΩ for PG, granule and deep SA cells) was not compensated. Voltages indicated in the paper were corrected for the junction potential (−15 mV).

### Drugs

6-nitro-7-sulfamoylbenzo[f]quinoxaline-2,3-dione (NBQX), D-2-Amino-5- phosphonopentanoic acid (D-AP5), 2-(3-carboxypropyl)-3-amino-6-(4 methoxyphenyl)pyridazinium bromide (GBZ), mecamylamine hydrochloride (MECA) and atropine were purchased from Abcam Biochemicals.

### Stereotaxic viral injections

3-6 weeks old dlx5/6-cre female mice were anesthetized with intraperitoneal injection of ketamine (20%), acepromazine (6%) and medetomidine (11.8%) mix (100µl/10g) and placed in a stereotaxic apparatus. Mice were craneotomized and a volume of 200-500 nl of AAV9.EF1a.DIO.hChR2(H134R).eYFP.WPRE.hGH or AAV9.EF1a.DIO.eYFP.WPRE.hGH (University of Pennsylvania Viral Vector Core) was stereotaxically injected with a Pneumatic Picopump PV 820, at 0.2 mm AP, −1.6 mm ML and −5.6 mm DV from bregma. After surgery, mice were injected with Metacam 100 (meloxicam) (5%) (100µl/10g), Antisedan (atipamezol) (2.5%) (100µl/10g), rehydrated with 1 ml of NaCl 0.9% and placed under a heating lamp. Mice recovered during 3-4 weeks after injection before anatomical or physiological experiments.

### Analysis

EPSCs, IPSCs or photocurrent amplitudes were measured as the peak of an average response computed from multiple sweeps. The decay of light-evoked IPSCs was most often best fitted with a double exponential with time t=0 at the peak of the current. Time constant values indicated in the text are weighted decay time constants calculated using the following equation: τ_w_=(τ_1_A_1_ + τ_2_A_2_)/(A_1_ + A_2_) where τ_1_ and τ_2_ are the fast and slow decay time constants and A_1_ and A_2_ are the equivalent amplitude weighting factors. The onset of EPSCs and IPSCs was measured at 5% of the first peak of the response. The latency of an ON-evoked EPSC was defined as the time interval between the beginning of the stimulation artifact and the onset of the EPSC. The latency of a light-evoked IPSC was defined as the time interval between the onset of light and the onset of IPSC. The jitter of evoked EPSCs and IPSCs was defined as the standard deviation (SD) of their latencies. To estimate the duration of an ON-evoked plurisynaptic excitatory response, individual EPSCs with amplitude >5 pA were automatically detected by the Axograph X software using a sliding template function. PSTH (peri stimulus time histograms) representing the cumulative number of EPSCs per 20 ms bin across several consecutive sweeps were constructed. We then determined the time needed after stimulation for the EPSC frequency to come back to baseline frequency + 2 SD during at least five consecutive bins. Baseline frequency was calculated over the 25 bins (i.e. 500 ms) preceding the stimulation.

### Statistics

Data are expressed as mean ± SD unless otherwise indicated in the text. We used the Student’s t-test to assess the statistical difference between paired sets of data with normal distribution, the non-parametric Wilcoxon-Mann-Whitney rank sum test to assess the statistical difference between unpaired sets of data, the Kruskal-Wallis test for data sets with more than two variables and p-values are reported.

### Immunohistochemistry

Adult mice (1-3 months) were deeply anesthetized with intraperitoneal injection of xylazine (6.3%) and ketamine (25%) and intracardially perfused with 4% paraformaldehyde (PFA). Brains were removed and kept in 4% PFA overnight. Slices were cut (50 µm thick) with a vibratome (Leica VT 1000S). Blocking steps were performed using a PBS solution containing 2% BSA or 5% goat serum and 0.3% Triton X-100 during 2h. Sections were incubated overnight at 4°C with primary antibodies: goat anti-choline acetyltransferase (ChAT; 1:500; Millipore Cat# AB144P, RRID:AB_2079751), mouse anti-parvalbumin (PV; 1:1000; Sigma-Aldrich Cat# P3088, RRID:AB_477329), rabbit anti-calretinin (CR; 1:1000; Swant Cat# 7697, RRID:AB_2721226) or mouse anti-calbindin D28K (CB; 1:1000; Sigma-Aldrich Cat# C9848, RRID:AB_476894). After three washes in PBS, sections were incubated 2 hours at room temperature with the corresponding secondary antibodies—Alexa fluor 633 conjugated goat anti-rabbit (1:500; ThermoFisher A-21070) and Alexa fluor 633 conjugated goat anti-mouse (1:500; ThermoFisher A-21050) or Alexa fluor 647 conjugated donkey anti-goat (1:200; ThermoFisher A-21447). After three washes, sections were mounted in Prolong Diamond Antifade Mountant (ThermoFisher). Images were taken using a Leica TCS SP5 II confocal microscope or an Axio Imager M2 for mosaic images. Immunostained and EYFP-expressing cells were manually counted using Cell counter plug-in on Fiji software.

## Results

### Selective expression of ChR2 in GABAergic neurons of the basal forebrain

We used an optogenetic approach to map the functional synaptic connections of basal forebrain GABAergic neurons onto olfactory bulb interneurons. We injected an adeno-associated virus encoding a Cre-inducible ChR2-EYFP fusion protein in the basal forebrain of dlx5/6-Cre mice, a transgenic line that expresses the Cre recombinase exclusively in GABAergic neurons in the forebrain (Monory *et al*., 2006)(Figure 1A). These injections produced a dense EYFP labeling of fibers in the olfactory bulb. Labeled fibers were particularly abundant in the granule cell layer and in the glomerular layer of the bulb, similarly to a previous study using a Cre-dependent viral approach in the GAD65-Cre mouse (Nunez-Parra *et al*., 2013). In the basal forebrain, expression of ChR2-EYFP was intense around the injection site but was limited to the membrane, making cell bodies difficult to distinguish. Hence, we injected the same virus expressing cytosolic EYFP to verify the specificity of the recombination in GABAergic neurons (Figure 1B, n=5 mice). EYFP was expressed in cell bodies of various sizes around the injection site. Some of these cells also expressed the calcium-binding proteins calretinin (CR, n=122 cells counted), calbindin (CB, n=54) or parvalbumin (PV, n=25), three classical markers of GABAergic neurons that label non-overlapping neural populations in the basal forebrain (Zaborszky *et al*., 2005). EYFP also colocalized with choline acetyltransferase (ChAT), the enzyme for acetylcholine synthesis, in about 1/3 of the ChAT(+) neurons counted (n=63/190 cells), consistent with the idea that cholinergic neurons in the basal forebrain are also GABAergic (Saunders *et al*., 2015).

**Figure 1:**
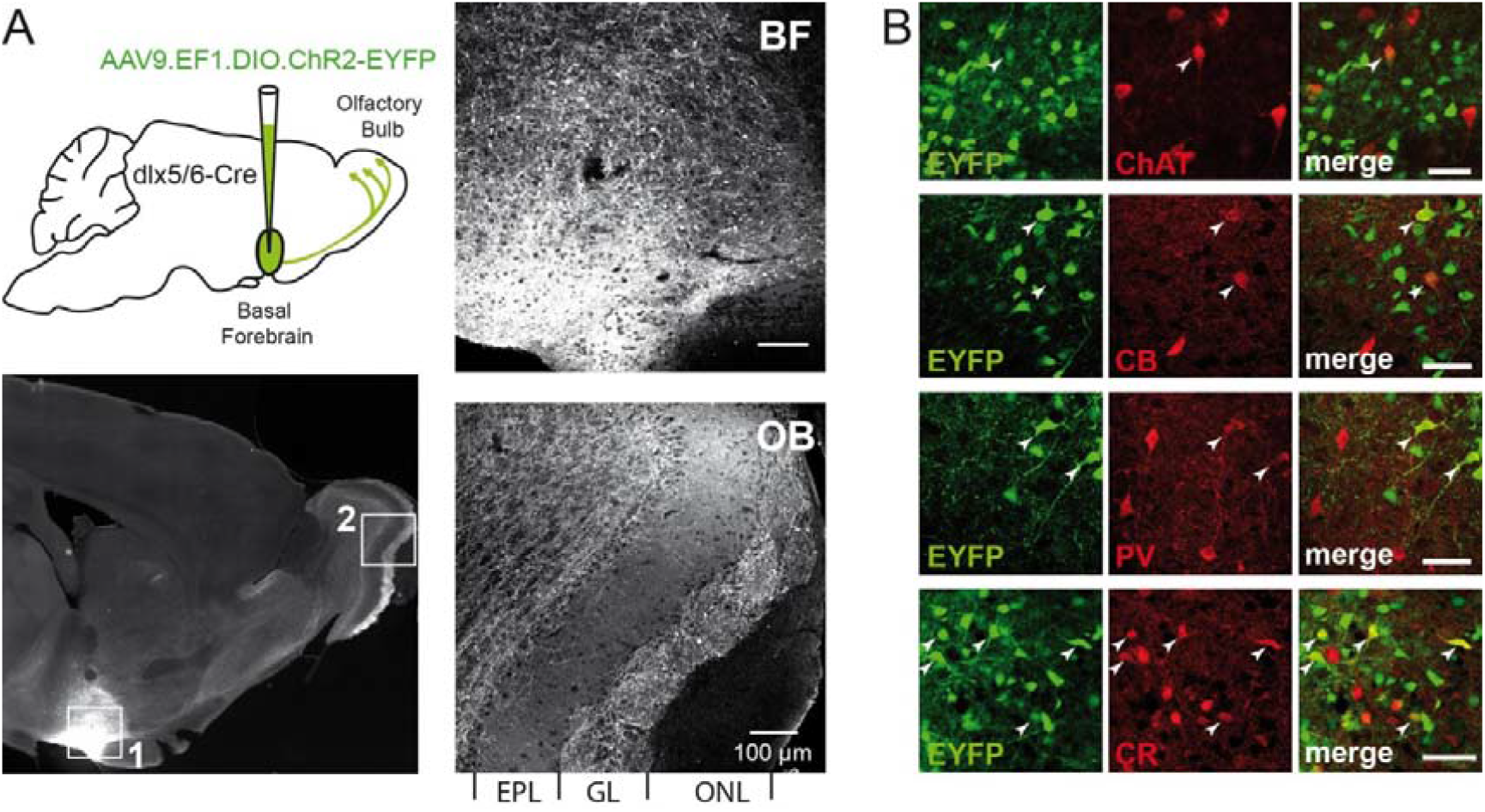
Conditional expression of ChR2 in various GABAergic neurons of the basal forebrain. **A,** Stereotaxic injection of a viral construct in the basal forebrain of dlx5/6- Cre mice was used for conditional expression of ChR2-EYFP in GABAergic neurons of the basal forebrain. Bottom left, sagittal section of a brain 4 weeks after the injection. The site of injection (BF, boxed area 1) and the olfactory bulb (OB, boxed area 2) are enlarged in the two other pictures. ONL: olfactory nerve layer, GL: glomerular layer, EPL: external plexiform layer. **B,** EYFP expression in the basal forebrain 4 weeks after injection of the same viral construct encoding cytosolic EYFP. Co-staining of EYFP (green) together with ChAT, CB, PV or CR (red). Some neurons (arrowheads), but not all, co-express the two markers. Scale bars 50 µm.

### Synaptic connections of basal forebrain axons in the olfactory bulb

We then tested the synaptic connections made by ChR2-expressing basal forebrain axons in the olfactory bulb using whole-cell voltage-clamp recordings in olfactory bulb slices. Consistent with previous reports (Nunez-Parra *et al*., 2013; Case *et al*., 2017), activating ChR2-expressing basal forebrain axons with brief (1-10 ms) flashes of blue light evoked outward inhibitory postsynaptic currents (IPSCs) in dSA cells (n=11/11, range 30-681 pA, mean 368±166 pA at Vh between −15 and −35 mV, Figure 2A) and in granule cells (n=10/12, range 31-464 pA, mean 148±124 pA at Vh=0 mV, Figure 2B). Light-evoked IPSCs had fast onset latencies (dSA: 1.6±0.35 ms, granule: 2.2±0.78 ms) and little jitter (dSA: 125±160 µs, granule: 274±170 µs), consistent with monosynaptic transmission. In 3 of the 11 dSA cells tested, the response had two components that reversed at different potentials. At Vh=-70 mV, the response was biphasic with an outward component that was blocked by the GABA_A_ receptor antagonist gabazine (GBZ, 5 µM), and an inward component that persisted in the presence of GBZ but was inhibited by the nicotinic receptor antagonist mecamylamine (MECA, 20 µM)(not shown). These dual responses are similar to those mediated by basal forebrain neurons that release both GABA and acetylcholine onto a specific subtype of dSA cells (Case *et al*. 2017) and were therefore not further studied.

**Figure 2.**
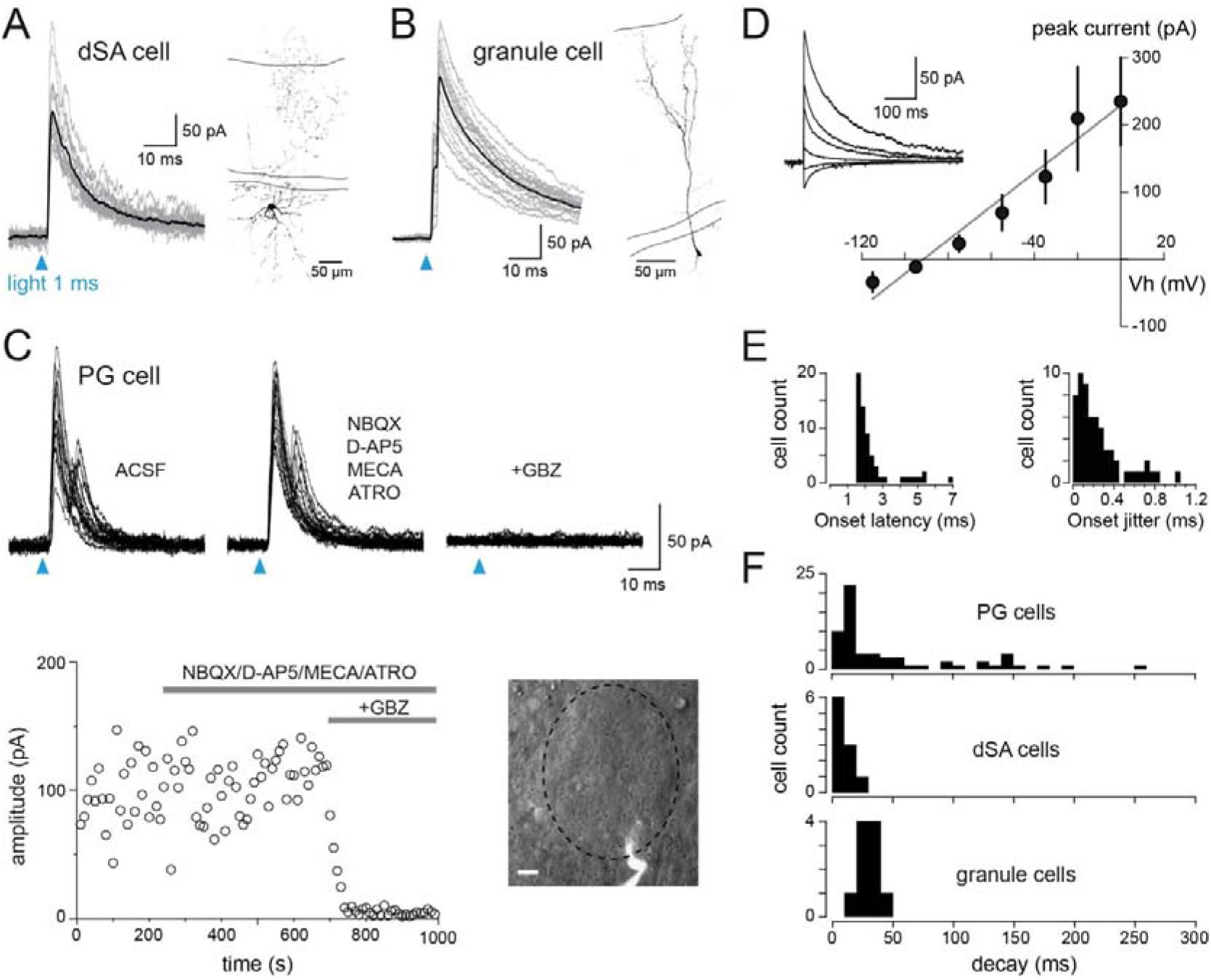
Synaptic connections of basal forebrain axons with various inhibitory interneurons of the olfactory bulb. **A,** Light-evoked consecutive ISPCs evoked in a deep SA cell recorded at Vh=-35 mV in the presence of AMPA and NMDA receptor blockers (10 µM NBQX + 50 µM D-AP5). The black trace is the average response. Light stimulation (1 ms) at blue arrow. The morphology of this neurobiotin-filled cell is shown. Solid lines delimit the glomerular layer (top) and the mitral cell layer (bottom). **B,** Same in a granule cell recorded at Vh=0 mV. **C,** Light-evoked IPSCs elicited in a PG cell (Vh=0 mV) in control condition, in the presence of NBQX (10 µM), D-AP5 (50 µM), mecamylamine (MECA, 20 µM) and atropine (ATRO, 10 µM) and after the addition of gabazine (5 µM, GBZ) to the cocktail. Several traces are superimposed for each condition. The graph shows the amplitude of consecutive responses in these different conditions. The image on the right is a snapshot of this PG cell during the recording, superimposed on the DIC image of the glomerulus delimited by a dashed line. Scale bar 10 µm. **D,** Average current-voltage relationship of the light-evoked IPSC in PG cells (n=12 cells, each plot is an average±sem). Inset, IPSCs from a representative PG cell recorded at different holding potentials. **E,** Distribution histograms of the onset latency (left) and onset jitter (right) of light-evoked IPSCs recorded in PG cells. **F,** Decay time constants distribution of light-evoked IPSCs recorded in PG, dSA and granule cells. PG and granule cells were recorded at Vh=0 mV, dSA cells at Vh comprised between −35 mV and −15 mV.

Most PG cells tested also responded to the photostimulation with an IPSC (n=73/94, range 6-974 pA, mean 231±218 pA at Vh=0 mV)(Figure 2C). In the majority of cells, light-evoked IPSCs had a short (<2 ms) onset latency that varied little across sweeps (jitter <200 µs), consistent with monosynaptic inputs (Figure 2E). However, in some cells, as in the case illustrated in Figure 2C, evoked IPSCs had longer and more variable onset latencies, displayed multiple peaks and had deflections during the rising phase. These complex responses resulting from summating asynchronous IPSCs could reflect spike asynchrony in presynaptic axons of different lengths. They could also arise from a multisynaptic pathway wherein light activation of the basal forebrain cholinergic afferents directly activate juxtaglomerular GABAergic neurons innervating PG cells or, indirectly, via the activation of mitral and tufted cells (Liu *et al*., 2015). To test this possibility, we blocked the glutamatergic transmission with the AMPA and NMDA receptor antagonists NBQX and D-AP5, respectively (n=14 PG cells), the cholinergic transmission with the nicotinic and muscarinic ACh receptor antagonists mecamylamine and atropine, respectively (n=6), or both pathways with a cocktail of these four drugs (n=4). These blockers had no effect on light-evoked IPSCs, even on complex ones (Figure 2C). Hence, our results are consistent with direct monosynaptic inputs of basal forebrain axons onto PG cells and confirm previous morphological evidence (Gracia-Llanes *et al*., 2010).

Evoked IPSCs recorded in ACSF or in the presence of acetylcholine and glutamate receptor blockers were then pooled for analysis. They were totally abolished by GBZ (n=11 PG cells, Figure 2C) and reversed at the expected equilibrium potential for chloride (ECl=-88 mV in our recording conditions, n=12 PG cells, Figure 2D). Light-evoked IPSCs had a uniformly fast time course in dSA cells (mean decay time constant 11.4±5.7 ms) and in granule cells (29±6.3 ms). In contrast, the duration of light-evoked IPSCs was highly variable in PG cells (decay time constant range 6-254 ms, mean 48.7±57.0 ms)(Figure 2F). However, light-responsive PG cells had diverse firing and synaptic properties, suggesting that the variable time course of their basal forebrain input may reflect PG cell diversity. To further validate this possibility, we classified the recorded PG cells in distinct subtypes based on their membrane properties, ON-evoked synaptic responses and light-evoked GABAergic synaptic inputs.

### Type 1 PG cells are not innervated by basal forebrain afferents

Type 1 PG cells constituted ∼10% of the PG cells recorded in dlx5/6;ChR2-EYFP mice (n=9 cells). These neurons fired multiple action potentials in response to a membrane supra-threshold step depolarization. Their discharge was regular (n=2), irregular (as in Figure 3A, n=3) or a burst of spikes (n=3). They responded to the electrical stimulation of OSN axons with a fast onset monosynaptic EPSC (mean latency 1.59±0.25 ms)(Figure 3B and 3E). The onset of this EPSC varied little across trials even at low intensities of stimulation (jitter 90±40 µs) whereas the response of type 2 PG cells had a longer onset latency (>2 ms) that increased and became more variable with stimulations of lower intensity (Figure 3E), as previously shown (Shao *et al*., 2009; Najac *et al*., 2015). ON stimulation also induced IPSCs in 8/9 type 1 PG cells voltage-clamped at Vh=0 mV, i.e. around the reversal potential for excitation (Figure 3B, 3D). This inhibitory response was composed of multiple asynchronous IPSCs, consistent with a plurisynaptic input generated within the glomerular network by other PG cells activated by the stimulation. However, none of the 9 type 1 PG cells tested responded to the photo stimulation of the basal forebrain fibers (Figure 3C). Thus, type 1 PG cells receive inhibitory inputs from neighboring PG cells but not from the basal forebrain.

**Figure 3:**
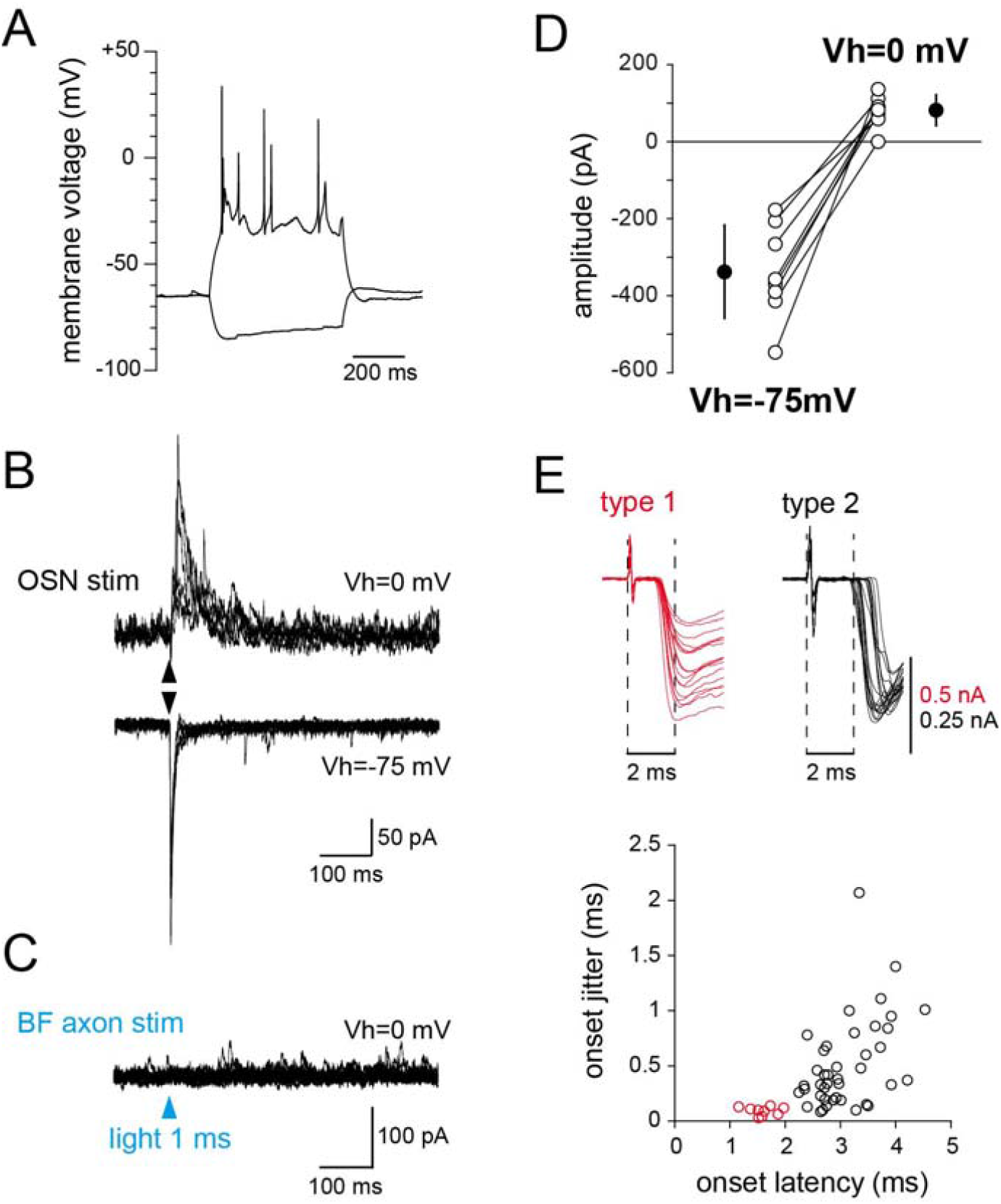
Type 1 PG cells are not contacted by basal forebrain projections but are inhibited by other PG cells. Membrane properties (**A**), ON-evoked synaptic responses (**B**) and responses to basal forebrain axons photo stimulation (**C**) of a type 1 PG cell recorded in a dlx5/6;ChR2-EYFP mouse. This cell responded to the stimulation of OSN axons with an inward fast monosynaptic EPSC at Vh=-75 mV and with outward plurisynaptic IPSCs at Vh=0 mV (**B**). In contrast, light stimulation of basal forebrain fibers did not evoke any response (**C**). Several traces are superimposed in each case. **D,** Summary plots for ON-evoked synaptic response amplitudes of all type 1 PG cells tested at two holding potentials. **E,** Classification of the recorded PG cells into type 1 and type 2 in dlx5/6;ChR2-EYFP mice. Top, early phase of ON-evoked EPSCs in a type 1 PG cell (red) and in a type 2 PG cell (black). The type 1 PG cell responded with a short latency EPSC, starting <2 ms after the beginning of the stimulation artifact and with little onset variability between trials. In contrast, responses from the type 2 PG cell had a delayed and more variable onset latency. The graph shows the onset latency vs. onset jitter of ON-evoked responses in PG cells classified as type 1 PG cells (red circles) and in PG cells classified as type 2 PG cells (black circles) in this study.

### Different classes of type 2 PG cells receive basal forebrain inputs with target-specific properties

Type 2 PG cells were largely predominant in our recordings in dlx5/6;ChR2-EYFP mice (n=68) and all PG cells responding to the flash of blue light with an IPSC were type 2 PG cells. We classified these type 2 PG cells in three subclasses, based on their membrane properties, ON-evoked synaptic responses and light-evoked basal forebrain inputs, as followed:

#### 1. CR-expressing type 2 PG cells

The most numerous cell type (n=29) showed the distinctive immature-like membrane and synaptic properties of CR-expressing PG cells (Fogli Iseppe *et al*., 2016; Benito *et al*., 2018), consistent with the predominance of this subtype within the entire population. These cells fired at most a single action potential or a spikelet and were passively depolarized by a suprathreshold current step (Figure 4A). In addition, these PG cells received few spontaneous EPSCs (not shown), had small ON-evoked excitatory inputs (Figure 4B, 4C) and, usually, a large electrical membrane resistance (Figure 4D). ON stimulations evoked small IPSCs or no IPSC at all in these cells clamped at Vh=0 mV (Figure 4B, 4C) as we previously reported in genetically-identified CR-expressing PG cells (Benito *et al*., 2018). In contrast, photo stimulation of basal forebrain fibers evoked a GBZ-sensitive IPSC in 27/29 cells (Figure 4E). These light-evoked IPSCs were often large (average amplitude 294±196 pA, range 6-741 pA) and always fast (average decay time constant 11.9±3.6 ms, range 6-21 ms)(Figure 4E). Thus, a fast basal forebrain-mediated GABAergic input is another attribute of CR-expressing PG cells.

**Figure 4:**
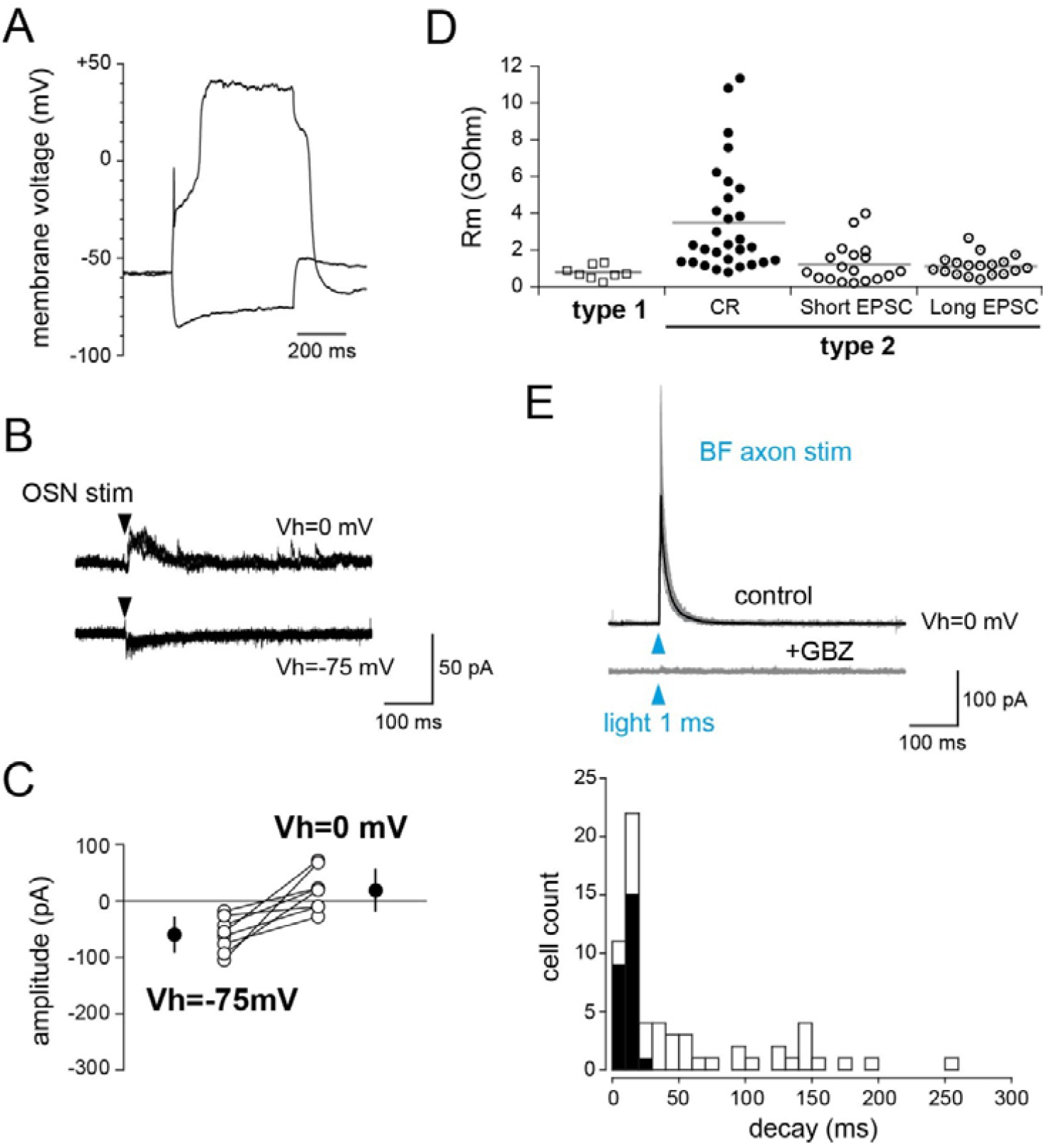
CR-expressing type 2 PG cells receive fast basal forebrain GABAergic inputs. Membrane properties (**A**) and ON-evoked synaptic responses (**B**) of a typical CR- expressing type 2 PG cell recorded in a dlx5/6;ChR2-EYFP mouse. This cell typically responded to a suprathreshold depolarization with a single spikelet followed by a passive membrane depolarization (**A**) and had small ON-evoked responses at Vh=-75 mV and at Vh=0 mV (**B**). **C,** summary plots for ON-evoked synaptic responses recorded at Vh=-75 mV and at Vh=0 mV for cells classified as CR-expressing PG cells. **D,** cells classified as CR-expressing type 2 PG cells also had larger electrical membrane resistances compared to PG cells classified in the other groups (horizontal bars show the averages, p<0.001 for comparison with each of the 3 other subgroups, Wilcoxon test). **E,** light stimulation of basal forebrain fibers evoked large and fast GBZ-sensitive IPSCs in CR-expressing PG cells. Traces are from the same cell as in A and B. The average IPSC (black trace) is superimposed on several consecutive responses recorded at Vh=0 mV in ACSF. Bottom, distribution histogram of the decay time constants of light-evoked IPSCs in cells classified as CR-expressing PG cells (black bars) superimposed on the distribution histogram for all the recorded PG cells (white bars).

#### 2. CB-like type 2 PG cells with short ON-evoked excitatory responses

The second group of type 2 PG cells (n=17) included cells that responded to ON stimulations with a short burst of asynchronous EPSCs that lasted only few tens of ms (average duration 41±27 ms)(Figure 5B). We have previously shown that this kind of response is observed in a heterogeneous subclass of PG cells that includes CB- expressing PG cells (Najac *et al*., 2015). Cells classified in this group were thus called CB-like PG cells. Their ON-evoked response is surprisingly much shorter than the firing of the presynaptic mitral and tufted cells, independent of the stimulation strength. However, the mechanism that limits glutamate release on these PG cells is still unknown. In the present study, the majority (n=9) responded to a depolarizing current step with a single spike preceding a plateau potential. However, some cells had a regular firing (n=3) and others responded with action potentials riding on a calcium spike (n=3). Photo stimulation of basal forebrain fibers evoked an IPSC in 12 of these 17 CB-like cells (Figure 5D). Light-evoked responses had a mean amplitude of 129±96 pA (range 24-324 pA) and a decay time constant ranging from 6.7 to 53.8 ms (mean 26±14 ms)(Figure 5D), significantly slower than in CR-like PG cells (p=0.0060, Wilcoxon test). In contrast, ON stimulation did not evoke IPSCs (Figure 5B, 5C), suggesting that these cells are not connected to other PG cells.

**Figure 5:**
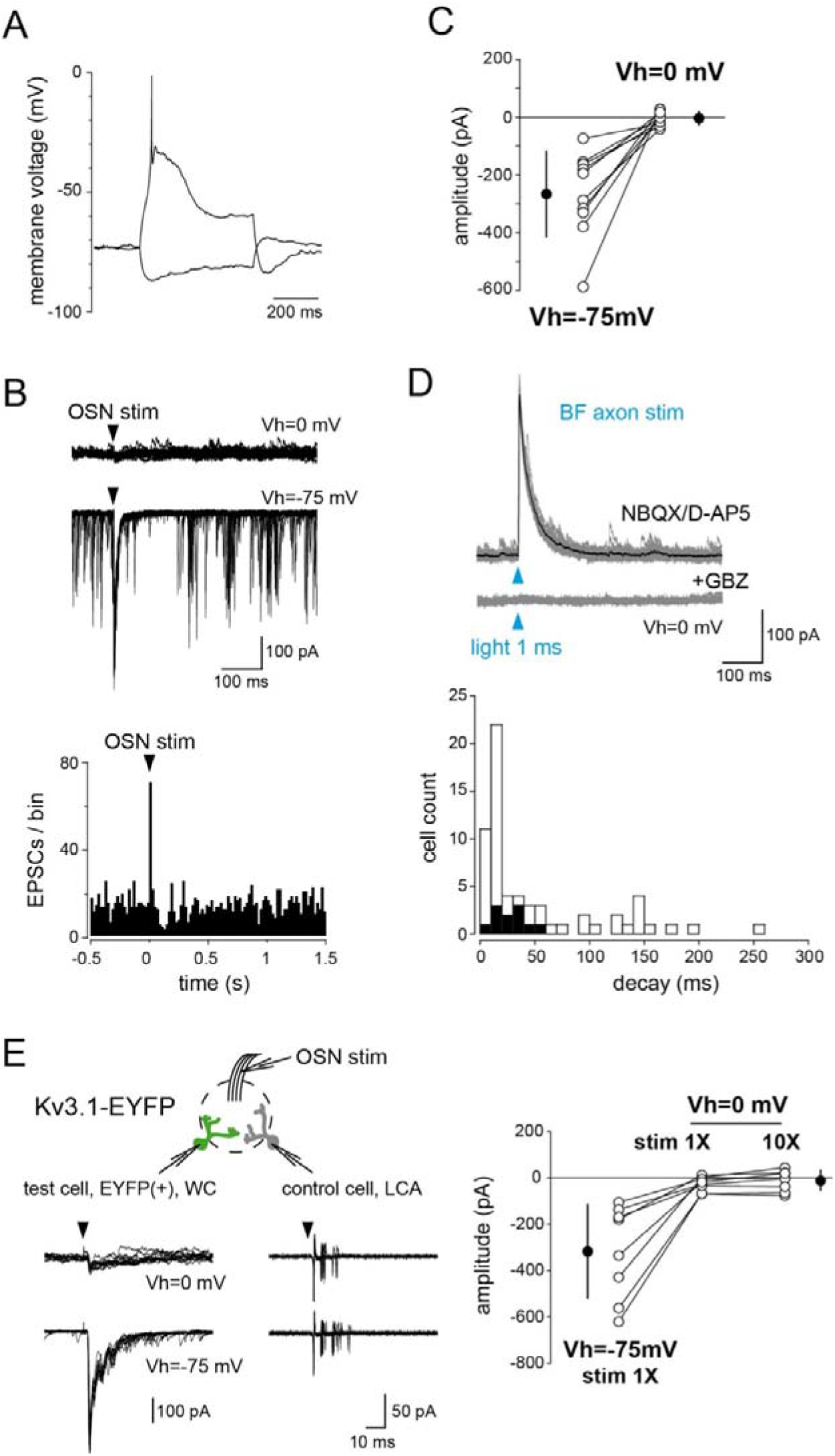
Type 2 PG cells with short ON-evoked excitatory response do not interact with other PG cells but receive inputs from the basal forebrain. Membrane properties (**A**) and ON-evoked synaptic responses (**B**) of a typical type 2 PG cell that responded to the stimulation of the OSN axons with a short burst of EPSCs in a dlx5/6;ChR2-EYFP mouse. The short ON-evoked excitatory response was the main criterion for classifying PG cells in this subclass. **C,** Summary plots (right) for ON-evoked synaptic responses recorded at Vh=-75 mV and at Vh=0 mV indicate that cells in this group also had no ON-evoked IPSC. **D,** IPSCs evoked by a light stimulation of the basal forebrain fibers. The average IPSC (black trace) is superimposed on several consecutive responses recorded at Vh=0 mV in the presence of NBQX and D-AP5. GBZ abolished the response. Traces are from the same cell as in A and B. Bottom, distribution histogram of the decay time constants of light-evoked IPSCs in PG cells classified in this subclass (black bars) superimposed on the distribution histogram for all the recorded PG cells (white bars). **E,** Paired recording of two PG cells projecting in the same glomerulus in the Kv3.1-EYFP transgenic mouse. The “test” EYFP(+) PG cell was recorded in the whole-cell configuration at Vh=0 mV to monitor IPSCs and at Vh=-75 mV to monitor EPSCs. The “control” PG cell was recorded in the loose cell-attached configuration to monitor its firing. OSNs were stimulated at an intensity inducing the firing of the control PG cell. The summary graph on the right shows that OSN stimulations did not evoke any outward IPSC in the test cell even when the stimulation was increased by a factor of 10.

To confirm the absence of connections with other PG cells, we used the Kv3.1-EYFP reporter mouse. A population of type 2 PG cells with short ON-evoked responses is labeled in this mouse, including CB-expressing cells (Najac *et al*., 2015). In these experiments, we recorded the whole-cell voltage-clamp response of EYFP(+) PG cells while simultaneously monitoring in loose cell-attached the discharge of a control PG cell projecting into the same glomerulus (either EYFP(+), n=4, or EYFP(-), n=5, Figure 5F). ON stimulations that efficiently drove the firing of the control PG cell induced a robust excitatory response in EYFP(+) PG cells clamped at Vh=-75 mV. This stimulation, however, did not evoke any IPSC when cells were clamped at Vh=0mV (n=9). In a subset of experiments, increasing the stimulation intensity by a factor of 10 also failed to evoke IPSCs (n=6). In contrast, an electrical stimulation in the glomerular layer at >200 µm away from the recorded PG cell evoked a reliable GBZ-sensitive IPSC in EYFP(+) PG cells (n=16, average amplitude 87±48 pA at Vh=0 mV; decay time constant: 30±13 ms, range 7.7-53.6 ms) that persisted in the presence of NBQX and D-AP5 (not shown). Our results suggest that this IPSC is mediated by basal forebrain axons ramifying in the glomerular layer, although the possibility that dSA cells also innervate these PG cells cannot be excluded. Thus, CB-like type 2 PG cells responding to the stimulation of the olfactory nerve with a short excitatory response do not interact locally with other PG cells but do receive a prominent GABAergic input from the basal forebrain.

#### 3. Type 2 PG cells with long-lasting ON-evoked excitatory responses

The third group of type 2 PG cells (n=18) does not correspond to any known genetically-identified PG cell subtype described so far. It is composed of cells that all responded to the injection of a depolarizing step with a regular firing (Figure 6A). On average, these cells fired up to 27±6 action potentials during a 500 ms-long suprathreshold step, a high frequency that distinguishes them from small TH(+) juxtaglomerular cells that fire at a lower rate (Galliano *et al*., 2018). In addition, they responded to the ON stimulation with a long-lasting barrage of EPSCs, often lasting several hundreds of milliseconds (average 1050±610 ms)(Figure 6B). These long responses also contrast with the short burst of EPSCs evoked in TH(+) juxtaglomerular cells (Kiyokage *et al*., 2010) but match well with the prolonged ON-evoked firing in mitral and tufted cells (Najac *et al*., 2015; Geramita & Urban, 2017). Moreover, 17/18 cells in this group responded to the photo stimulation of the basal forebrain axons with a prolonged IPSC (IPSC amplitude 193±247 pA, range 11-850 pA; average decay time constant 116±63 ms, range 40-254 ms, p<0.0001 for comparison with both CR- like and CB-like PG cells, Wilcoxon test)(Figure 6D). These basal forebrain-mediated IPSCs were much larger than the ON-evoked IPSCs that were eventually observed in 6/11 PG cells (Figure 6C). Thus, long-lasting basal forebrain-mediated IPSCs are found in a specific and homogeneous subclass of regularly firing PG cells that responds to an ON input with a remarkably prolonged barrage of EPSCs.

**Figure 6:**
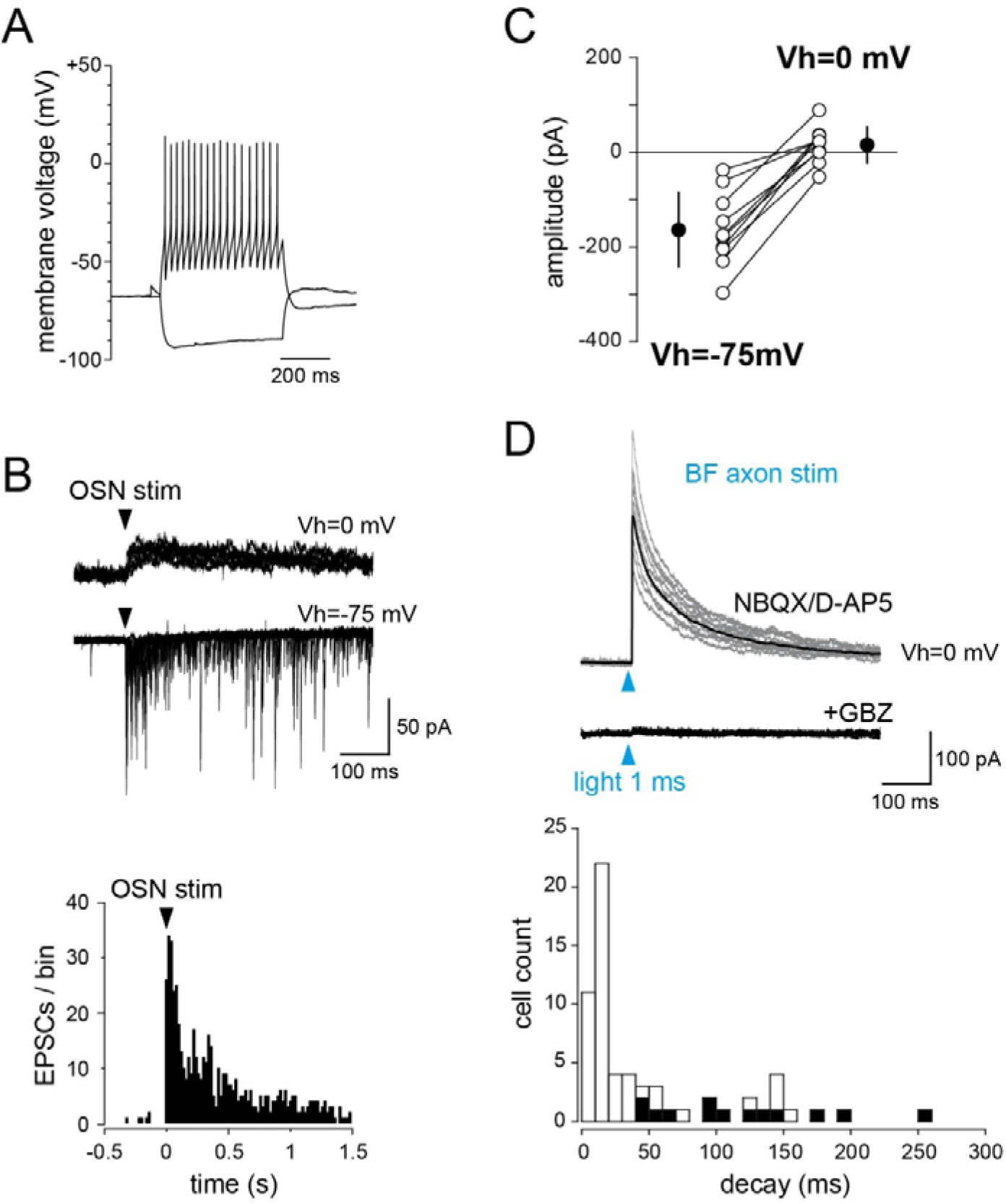
Type 2 PG cells with long-lasting ON-evoked excitatory response receive slow GABAergic inputs from the basal forebrain. Membrane properties (**A**) and ON- evoked synaptic responses of a typical type 2 PG cell that responded to the stimulation of the OSN axons with a long-lasting burst of EPSCs in a dlx5/6;ChR2-EYFP mouse. **A,** Like all the cells included in this group, this PG cell responded to a step depolarization with a regular firing. **B,** ON-evoked responses recorded at two holding potentials and PSTH of the excitatory response (bottom). **C,** Summary plots for ON-evoked synaptic responses recorded at Vh=-75 mV and at Vh=0 mV for cells included in this group. **D,** IPSCs evoked by a light stimulation of the basal forebrain fibers. The average IPSC (black trace) is superimposed on several consecutive responses recorded at Vh=0 mV in the presence of NBQX and D-AP5. GBZ abolished the response. Same cell as in A and B. Bottom, distribution histogram of the decay time constants of light-evoked IPSCs in PG cells classified in this subclass (black bars) superimposed on the distribution histogram for all the recorded PG cells (white bars). Cells included in this group had slow IPSCs.

Finally, 21 of the recorded PG cells, that either had an incomplete characterization (n=17) or functional properties that did not fit in any of the 4 previously defined subgroups (n=4), were not classified. 17 of these cells responded to the photo stimulation with an IPSC.

### Diversity of basal forebrain afference

Our data so far indicate that the time course of the basal forebrain synaptic inputs depends on the PG cell subtype they target. To start gaining insight into whether these distinct postsynaptic PG neurons are contacted by different presynaptic fibers, we compared the short-term plasticity at these synapses. We applied a train of 5 blue light pulses at 20 Hz. This photo stimulation evoked IPSCs that depressed at different degrees in the three subclasses of type 2 PG cells as quantified by the paired-pulse ratio of the second IPSC amplitude relative to the first (Kruskal-Wallis test, H=11.19, p=0.0037)(Figure 7). In particular, the paired-pulse depression in CR-like PG cells (0.73±0.13, n=11) was less pronounced than in CB-like PG cells (0.46±0.16, n=7, p=0.0012, Wilcoxon test) and than in PG cells with long-lasting ON-evoked responses (0.56±0.16, n=8, p=0.020, Wilcoxon test). The paired-pulse ratio was not different in these two last groups (p=0.28, Wilcoxon test) but failures of transmission were frequent in CB-like PG cells (seen in 5/7 cells, Figure 7B) whereas they were never observed in PG cells with long-lasting ON-evoked responses. Together, these data provide evidence that basal forebrain inputs may be mediated by specific afferent fibers on each subclass of olfactory bulb PG cells.

**Figure 7:**
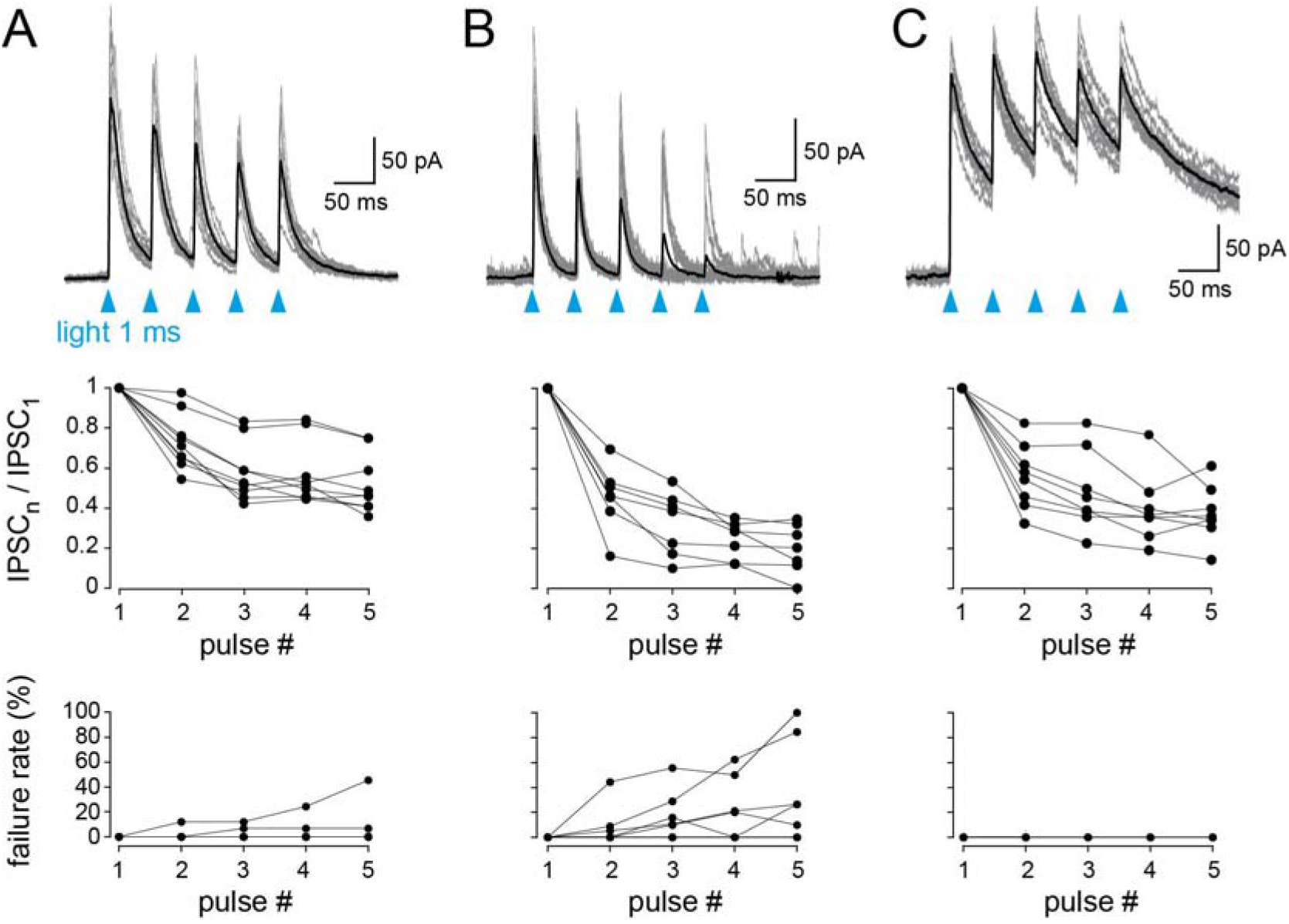
Basal forebrain GABAergic inputs have different presynaptic properties depending on the postsynaptic PG cell subtype. **A-C, Top row**, light-evoked IPSCs in three PG cells representative of the three subclasses of type 2 PG cells recorded in dlx5/6;ChR2-EYFP mice (**A:** CR-expressing PG cells; **B:** PG cells with short ON-evoked excitatory responses and **C:** regularly firing PG cells with long-lasting ON-evoked responses). Each cell was stimulated with 5 flashes of light at 20 Hz. 10-12 consecutive responses are superimposed for each cell, the black trace is the average response. **Middle row**, amplitudes of the n^th^ light-evoked IPSC relative to the normalized amplitude of the first IPSC recorded in PG cells classified in the 3 subgroups of the corresponding column. Lines connect plots from the same cells. **Bottom row**, failure rate for each of the five consecutive stimuli and for each cell over multiple trials. Failures were never observed in the third group.

## Discussion

Most efforts in understanding the function of inhibition in the olfactory bulb have focused on the local activation mechanisms of the diverse interneuron populations and on the impact of their activity in shaping the olfactory bulb output. In comparison, GABAergic circuits that control the activity of olfactory bulb interneurons have received little attention. We show here that long-range GABAergic projections from the basal forebrain make prominent and specific connections on most classes of interneurons in the olfactory bulb with a previously unsuspected level of synaptic specificity. These data provide additional criteria for PG cells classification and emphasize that each PG cell subtype belongs to a specific synaptic circuit, an important step towards understanding the functional implications of their diversity. They also reinforce the idea that the basal forebrain is a major modulator of olfactory bulb circuits whose influence is still partially understood.

### Basal forebrain projections to the olfactory bulb

The basal forebrain contains several nuclei without clear delineations that innervate the cerebral cortex, limbic areas, hippocampus, thalamic structures and the olfactory bulb with cholinergic and GABAergic projections. Centrifugal projections to the olfactory bulb originate principally from the HDB and MCPO. Cholinergic axons are found essentially in the glomerular layer and in the internal plexiform layer of the bulb and, moderately, in the granule cell layer (Macrides *et al*., 1981; Zaborszky *et al*., 1986; Salcedo *et al*., 2011; Case *et al*., 2017; Hamamoto *et al*., 2017). GABAergic projections are even more massive and innervate all layers of the bulb (Zaborszky *et al*., 1986; Gracia-Llanes *et al*., 2010; Niedworok *et al*., 2012; Nunez-Parra *et al*., 2013). Cholinergic and GABAergic neurons are not clearly segregated in the basal forebrain (Zaborszky *et al*., 1986), interact between each other (Yang *et al*., 2014) and nearly all cholinergic neurons express the molecular machinery to co-transmit GABA (Saunders *et al*., 2015). These two centrifugal pathways are thus intimately coupled and attempts to evaluate their influence independently have to be interpreted with caution. Nevertheless, several reports demonstrate that ACh modulates diverse bulbar neurons at different timescales and via multiple pre-and postsynaptic muscarinic and nicotinic receptors (Castillo *et al*., 1999; Ghatpande & Gelperin, 2009; Devore *et al*., 2014; Liu *et al*., 2015; Smith *et al*., 2015). Optogenetic manipulation of basal forebrain cholinergic neurons modulates odor detection, discrimination and olfactory learning (Ma & Luo, 2012; Rothermel *et al*., 2014). Our results indicate that basal forebrain GABAergic neurons also have multiple targets in the bulb. Not surprisingly, pharmacological inactivation of HDB/MCPO GABAergic neurons using a DREADD/CNO paradigm impairs olfactory sensitivity and odor discrimination (Nunez-Parra *et al*., 2013). The basal forebrain therefore stands as a major modulator of olfactory bulb circuits but understanding its global impact on olfactory bulb processing may first require a careful investigation of its action on each targeted neuronal population.

### GABAergic synaptic connections discriminate different classes of PG cells

We closely examined the GABAergic inputs of olfactory bulb PG cells. The results demonstrate that GABAergic connections are an additional criterion that helps classifying diverse PG cell subtypes. First, type 1 and type 2 PG cells can be distinguished on the basis of their inhibitory input. Basal forebrain fibers contact different subtypes of type 2 PG cells but not type 1 PG cells. Moreover, type 2 PG cells have little synaptic interactions with other PG cells whereas type 1 PG cells do. Previous studies revealed that GABAergic lateral interactions between PG cells projecting within the same glomerulus are mediated, at least in part, by GABA spillover (Smith & Jahr, 2002; Murphy *et al*., 2005). Our results further suggest that more conventional PG-PG cells synaptic interactions may be restricted to type 1 PG cells.

Second, we show that basal forebrain axons produce IPSCs with distinct time courses and short term plasticities in three subgroups of type 2 PG cells that also differ on the basis of their ON-evoked excitatory response and firing properties. IPSCs with diverse durations likely reflect different subunit compositions of postsynaptic GABAA receptors. For instance, a subpopulation of PG cells that does not overlap with CB-, CR- and TH-expressing cells selectively expresses the alpha5 subunit of the GABA_A_ receptor (Panzanelli *et al*., 2007). Expression of this subunit correlates in some neurons with slow IPSCs (Pangratz-Fuehrer *et al*., 2016) suggesting that the alpha5 subunit may be a molecular marker of the type 2 PG cells with regular firing and a long-lasting ON- evoked excitatory response.

Postsynaptic GABAA receptors with different kinetics contribute in shaping short term depression. However, our analysis of paired pulse ratio and failure rate suggest that the different degrees at which basal forebrain IPSCs depressed in PG cells also reflect presynaptic diversity. Thus, distinct subsets of chemically-defined basal forebrain GABAergic neurons may control specific PG cell subclasses. Consistent with this high level of centrifugal afferent selectivity, a recent report demonstrates that a specific subset of GABA/ACh basal forebrain neurons preferentially projects in the internal plexiform layer of the olfactory bulb where it selectively connects with one specific subtype of dSA cells (Case *et al*., 2017). Thus, addressing the synaptic complexity of basal forebrain afferents in the glomerular layer of the olfactory bulb may necessitate the use of transgenic mice allowing the manipulation of selective genetically-identified population of neurons.

### Are there other classes of PG cells?

We have defined four subclasses of PG cells based on functional parameters. However, we limited our characterization to a subset of electrophysiological properties that, in our recording conditions, could be unambiguously established. A full characterization that correlates membrane and synaptic properties with morphological and molecular features would most likely define additional subclasses of PG cells. For instance, type 2 PG cells with short-lasting ON-evoked excitatory responses have heterogeneous firing patterns and include CB-expressing and CB-lacking PG cells (Najac *et al*., 2015). In addition, some TH(+) juxtaglomerular neurons may constitute a subclass of PG cells. These GABAergic/dopaminergic neurons constitute 20% of the juxtaglomerular GABAergic cells (Panzanelli *et al*., 2007; Parrish-Aungst *et al*., 2007). They are often collectively called superficial SA cells although only a small fraction of them corresponds to the classical definition of superficial SA cells i.e. large TH(+) cells with an axon and dendrites that widely ramify in the glomerular layer (Kosaka & Kosaka, 2008; Kiyokage *et al*., 2010; Galliano *et al*., 2018). The majority though is anaxonic and resembles the classical PG cells, with a small cell body and processes that locally ramify, sometime within a single glomerulus (Kosaka & Kosaka, 2008; Galliano *et al*., 2018). We cannot exclude that small TH(+) cells have been included in our analysis but they would constitute at most 1 out of 5 of our randomly recorded PG cells. Moreover, their electrophysiological properties do not match well with any of the four subclasses we defined. Small TH(+) juxtaglomerular cells fire spikes in a regular manner but at low rate (Pignatelli *et al*., 2005; Puopolo *et al*., 2005; Galliano *et al*., 2018) and a majority responds to the ON stimulation with a short polysynaptic response (Kiyokage *et al*., 2010). Thus, our study does not address whether TH- expressing juxtaglomerular neurons also receive basal forebrain inputs and future experiments done in genetically identified TH(+) PG cells are needed to address this important question.

### Role of basal forebrain GABAergic afferents in olfactory bulb odor processing

We show the connections of basal forebrain GABA fibers with PG and granule cells that, together, constitute the main populations of interneurons directly regulating the activity of olfactory bulb output neurons. Deep SA cells also provide inhibitory input on PG and granule cells (Eyre *et al*., 2008; Burton *et al*., 2017) and are themselves modulated by basal forebrain inputs. This diversity of olfactory bulb interneurons under the control of basal forebrain fibers makes it challenging to predict the global functional implications of this GABAergic innervation on the olfactory bulb output. In the simplest way, inhibition of olfactory bulb interneurons may induce a disinhibition of mitral and tufted cells and thus facilitate the olfactory bulb output. However, GABA may be excitatory on some PG cells (Parsa *et al*., 2015). Moreover, co-transmission of GABA and ACh on, at least, some interneurons (Case *et al*., 2017), also complicates this view. Moreover, as olfactory bulb processing is fundamentally rhythmic and tightly coupled to respiration, the physiological impact of basal forebrain centrifugal inputs may depend on their temporal dynamics along a respiration cycle. A synchronized activity of these centrifugal afferents may reset the activity of olfactory bulb interneurons before a new respiration cycle. But how synchronized are they? Recent data indicate that some non-cholinergic neurons in the HDB have their activity correlated with attention whereas the activity of cholinergic neurons is correlated with primary reinforcers and outcome expectations (Hangya *et al*., 2015). Centrifugal inputs from the basal forebrain could therefore contextually modulate the olfactory bulb activity as a function of the internal state.

Interestingly, basal forebrain GABAergic fibers and axonal feedback from the olfactory cortex target the same populations of interneurons in the bulb. Thus, axons from pyramidal cells of the piriform cortex or the anterior olfactory nucleus directly excite olfactory bulb interneurons and facilitate mitral cell inhibition (Boyd *et al*., 2012; Markopoulos *et al*., 2012), i.e. the exact opposite of what HDB GABAergic fibers might do. Pyramidal cells of the olfactory cortex innervate HDB neurons as well (Paolini & McKenzie, 1997; Linster & Hasselmo, 2000) (Do *et al*., 2016). Centrifugal afferents from the basal forebrain could have complex dynamics, similar as cortical feedback that exerts spatially diffuse, temporally complex and non-uniform actions on olfactory bulb interneurons in response to odors (Boyd *et al*., 2015; Otazu *et al*., 2015). Thus, the net impact of cortical feedback and basal forebrain fiber inputs on olfactory bulb interneurons, balancing or opposite actions, may depend on their relative timing in different behavioral context.

## Acknowledgements

This work was supported by the Centre National pour la Recherche Scientifique, the Université de Strasbourg and the Agence Nationale pour la Recherche (Grant ANR-12- JSV4-006-01). ASD was funded by a fellowship from the Ministère de la Recherche and by a fellowship from the Fonds Paul Mandel pour les Recherches en Neuroscience. We thank Ipek Yalcin (Institut des Neurosciences Cellulaires et Intégratives, Strasbourg) and Claire Gaveriaux-Ruff (Institut de Génétique et de Biologie Moléculaire et Cellulaire, Strasbourg) for the kind gift of the dlx5/6-Cre mice and Thomas Knöpfel (Imperial College, London) for the kind gift of the Kv3.1-EYFP mice. We thank Sophie Reibel-Foisset and the animal facility Chronobiotron (UMS 3415 Centre National pour la Recherche Scientifique and Université de Strasbourg), the Plateforme Imagerie In Vitro-Neuropôle-Strasbourg, and Aline Huber for their technical assistance. We thank Jean Christophe Cassel (Laboratoire de Neurosciences Cognitives et Adaptatives, Strasbourg) Nuria Benito, Philippe Isope, Matilde Cordero-Erausquin, Bernard Poulain, and members of the lab for their constructive comments throughout the project.

## Competing interests

The authors declare no competing financial interests.

Authors contributions
ASD and DDSJ did the experiments, MN contributed essential preliminary data, DDSJ wrote the paper. All authors edited the manuscript.

